# Invasion histories reveal most North American introduced plants have not yet reached climatic stasis

**DOI:** 10.64898/2026.03.05.709936

**Authors:** Maisy Roach-Krajewski, Tyler W. Smith, Heather M. Kharouba

## Abstract

**Aim:** Analysis of species distributions often rests on the assumption of environmental equilibrium. That is, the distribution of a species (as documented by observation records) captures the full range of environmental conditions under which that species can maintain viable populations. Despite the centrality of this assumption to a variety of biogeographic questions, it is rarely empirically tested. This is particularly critical for recently introduced invasive species that are characterized by rapid expansion in their introduced range, often coupled with a niche shift relative to their native distribution. Defining equilibrium under these dynamic conditions is difficult. We developed the concept of environmental stasis as a more tractable proxy for equilibrium. In the context of species invasions, we define stasis as a prolonged period without an increase in the environmental conditions occupied by a species.

**Location:** North America

**Time Period:** 1614 to 2020.

**Major Taxa Studied:** Invasive plants

**Methods:** We applied the metric of climatic stasis to a suite of 258 invasive plant species in North America. We categorized their invasion trajectories into three classes (linear, two- and three-phase) based on theoretical expectations and then assessed how many had demonstrated environmental (climatic) stasis over a period of at least thirty years.

**Results:** More than 80% of the species were best fit by two- or three-phase models, indicating a declining rate of expansion. Climatic stasis was only documented for 44% of the species. In contrast, 85% of the species were in climatic stasis in their native ranges. The time to reach stasis ranged from 30 to 145 years (mean 90), and species at stasis in their invaded range occupied 97% of the climatic space they occupied in their native range.

**Main Conclusions:** This assessment provides valuable insight into the unrealized threat posed by the majority of invasive plants that have not yet reached stasis, as well as identifying which species can be most appropriately evaluated by methods that depend on the equilibrium assumption. Our work also demonstrates the useful perspective provided by the environmental stasis concept, which enables empirical quantification of one of the key aspects of equilibrium.

## Introduction

As climate change accelerates the spread of introduced plants across North America (Bradley et al., 2024), it has become increasingly important to understand and anticipate their expansion patterns within new regions. Because early detection and prevention are more effective than eradication after establishment (Jarnevich et al., 2023; Rockwell-Postel et al., 2020), managers would like to know where invasions are most likely to occur to prioritize those areas.

The range expansion of introduced species has been frequently predicted using species distribution models (SDMs), a widely used approach in ecology and conservation biology. SDMs relate georeferenced occurrence records (presence-only or presence/absence) to a set of environmental variables (e.g., climate, land cover) to predict habitat suitability across geographic space (Václavík & Meentemeyer, 2012). Despite the widespread use of SDMs, producing reliable predictions for the range expansion of introduced species remains challenging. One of the fundamental assumptions of the approach is often not supported: that the species is at environmental equilibrium, i.e., that it occupies all suitable environments, while unsuitable environments remain unoccupied (Araújo et al., 2005; Gallien et al., 2012). However, introduced species may be invasive, undergoing rapid and aggressive range expansion (Richardson et al., 2000; Pysek et al., 2020), often coupled with a niche shift relative to their native range (Wiens et al. 2005; Atwater et al. 2018),and may only near equilibrium in the final stages of invasion (Václavík & Meentemeyer, 2012; Bradley et al., 2015; Foster et al., 2022). Thus, SDMs based on data collected before this point may not produce accurate predictions (Gallien et al., 2012; Briscoe Runquist et al., 2019).

Despite the potentially wide-reaching implications, relatively few studies have attempted to quantify environmental equilibrium at broad scales. When they do, equilibrium is typically evaluated from a snapshot in time (e.g., Gallien et al. 2012; Bradley et al. 2015; Pili et al., 2020; Goncalves et al., 2022) and based on the comparison between a species’ current (i.e., *realized*) and *potential* distribution where any mismatch indicates the species is not at equilibrium. The potential distribution is often based on the species’ niche in its native range. These native niche requirements are then projected onto the invaded range to determine the potential suitable area the species will occupy at equilibrium (Petitpierre et al. 2017).

There are two main issues with this approach. First, it assumes that the species is at equilibrium in its native range, which may not be true (Early & Sax, 2014). If a species is not at equilibrium in its native range, projecting its distribution to the invaded range will underestimate its fundamental niche. Conversely, there may be records from areas in the native range where the species no longer occurs (e.g., due to shifting climate). These data could overestimate its fundamental niche. Second, this approach assumes that the equilibrium conditions in the invaded and native range are the same (niche conservatism; Wisz et al., 2013, Mainali et al. 2015). However, differing selection pressures in the invaded range may lead to local adaptation of the invading population, leading to a shift in its fundamental niche (Wiens et al. 2005). As such, the potential distribution in the native range may not truly reflect the potential niche of the species in its invaded range.

The problem may be more tractable if we shift our goal from quantifying equilibrium to assessing one of the most consequential symptoms of meeting the assumption: a species that is currently at environmental equilibrium will occupy the same environmental space over ecological time scales (i.e., years to decades, hereafter ‘environmental stasis’). In contrast, the environmental space of a species not at equilibrium will shift as it expands into previously unoccupied, but environmentally suitable areas. While environmental stasis is not sufficient to establish that a species is at environmental equilibrium, it is a necessary condition for it. Conversely, a species not in environmental stasis is demonstrably not at equilibrium.

Here we show that an assessment of current environmental stasis can be evaluated using historical observation records. Theory predicts a species should expand its distribution towards equilibrium as it proceeds through the stages of invasion (Theoharides & Dukes, 2007; Václavík & Meentemeyer, 2012, Foster et al., 2022; Figure 1). Invasions often follow a three-phase pattern: initial slow growth (the lag phase), followed by rapid spread, and finally equilibrium (Figure 1; Hui and Richardson 2017). During establishment, the environment occupied by an invasive species represents a small subset of the total available suitable environment (i.e., its equilibrium distribution; Radosevich et al., 2003). Thus, the species is furthest from equilibrium at this stage, and the predictions from SDMs are expected to be least reliable (Briscoe Runquist et al., 2019; Foster et al., 2022). After the species has become established (i.e., developed self-sustaining local populations), the invading species rapidly expands into previously unoccupied suitable climates. The species reaches environmental stasis, no longer expanding in environmental space once it has reached its equilibrium distribution. Therefore, to know *if* or *when* a species has reached equilibrium, a temporal context is essential (Foster et al. 2022).

**Figure 1.**
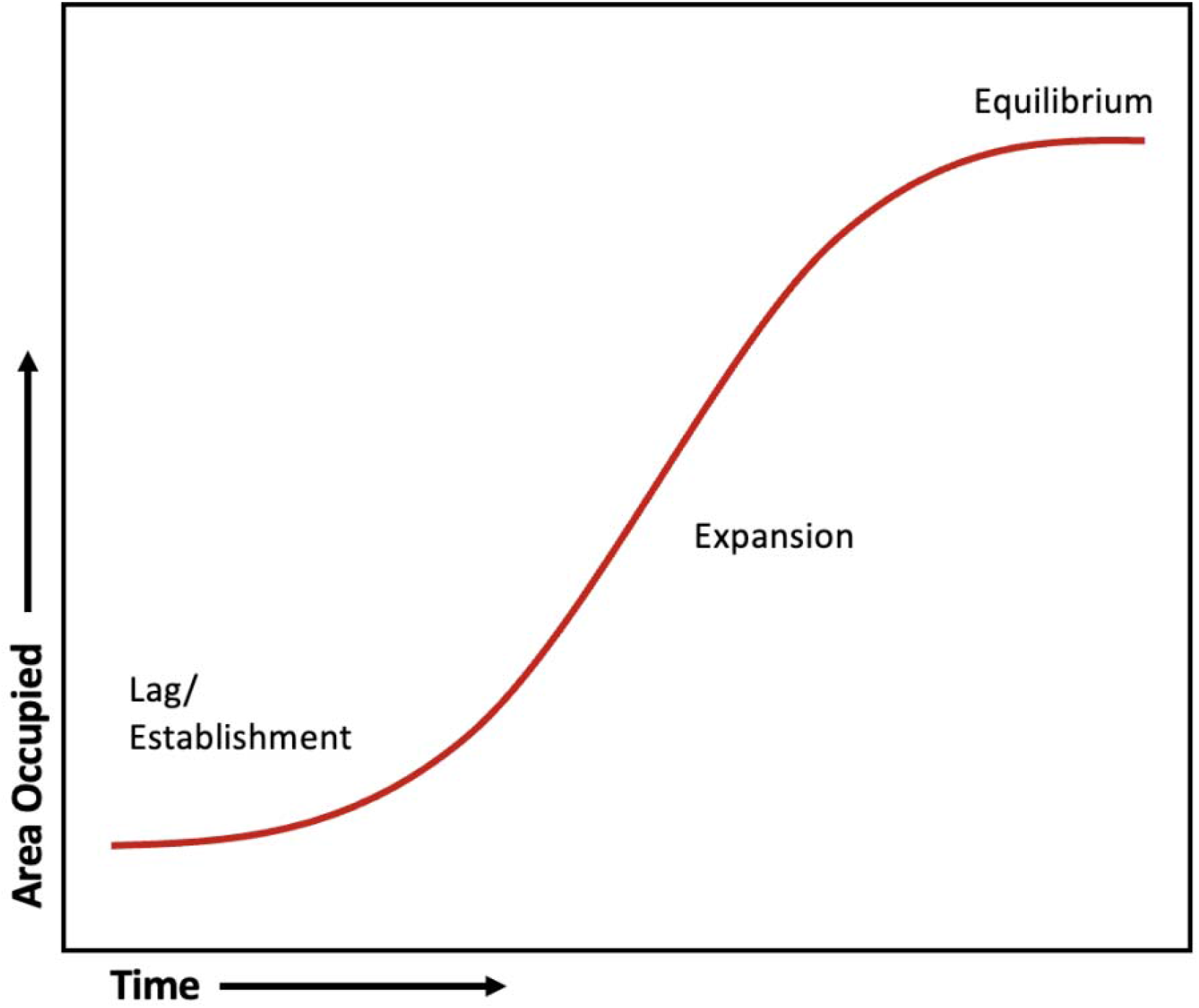
Theoretical curve depicting the three phases of the invasion process: lag/establishment phase, expansion phase, and equilibrium phase.

Quantifying patterns of expansion for a large dataset of invasive species using historical data is not a new concept. However, past studies – particularly those at macro-scales – have looked at patterns of expansion in *geographic* space only (i.e., how occurrences accumulate in physical space; e.g., Mosena et al., 2018; Aiello-Lammens, 2020; Osunkoya et al., 2021). This approach does not allow us to accurately distinguish between expansion into novel (i.e., not occupied in the native range) or similar (i.e., occupied in the native range) environments in the introduced range, thus constraining our ability to predict where expansions might occur in the future.

In this study, we determine if, and when, 258 introduced plant species in North America reached climatic stasis. To do so, we analyze their spread in climate space in the invaded range over the invasion history in North America. At 5-year time intervals, we quantify the climate space that a species occupies in its invaded range relative to that occupied in its native range. As an invasive species progresses through its invasion, more climate space should become occupied over time (Figure 1), thus occupying a growing subset of the climate space known to be suitable based on its native distribution. Using this metric, we first quantify the pattern of spread into climate space over time and test whether invasive plants typically follow a three-phase pattern. Second, we determine if a species has reached climatic stasis in its introduced range: a prolonged period (at least 30 years) without spread. Third, we assess whether species have reached stasis in their native range, and for species at stasis in both ranges we compare the environments occupied in each. To the best of our knowledge, this study represents one of the first to use a temporal approach to describe the equilibrium status of a large group of introduced species.

## Methods

### Data collection

We compiled a database of georeferenced specimen records from the Global Biodiversity Information Facility (GBIF; DOI: 10.15468/DD.TYMJWM) for non-native, invasive plant species in North America based on the list of species compiled by Atwater et al. (2018), from which we selected terrestrial tree, shrub and forb species. To ensure a long enough time-series and reasonable sample size for the analysis, we eliminated species without *i)* at least one occurrence in North America prior to 1990, and *ii)* at least 20 total occurrence records in each of their native range and introduced range. This resulted in a final dataset including 258 species from 61 plant families with five native ranges (Table S1).

To define the climate space occupied by each species in their native and invaded (i.e., North American) range, we imported climate data from WorldClim version 2.1 for 1970-2000 (Fick and Hijmans 2017) at a resolution of 10 km^2^. This included 19 variables derived from monthly temperature and rainfall values. To represent only the climatic combination present in these ranges and to improve the resolution of the spatial analysis, the climate rasters were cropped into six separate geographic extents: one to match the invaded (North American) range, and one matched to each of the five different native ranges. The climate space for each species was defined by the first two axes of a principal components analysis (PCA) combining the climate data from its native range and the invaded North American range, divided into a 100 x 100 = 10,000 cell grid. Thus, we used five different climate grids to capture the environments occupied by species from each of the five different native ranges.

### Quantifying spread

We modelled the spread in climate space for each species at 5-year intervals throughout their invasion history in North America using the *ecospat* package in R (Di Cola et al. 2017). We compared the density of occurrence records in the invaded range at the end of each time period with the current distribution (as of 2020) in its native range. Ecospat partitions invasion data into three categories relative to a reference native distribution: conditions occupied in both the invasive and native ranges are classified as ‘stable’, conditions only occupied in the invaded range are ‘expansion’, and conditions occupied only in the native range are ‘unfilling’. Preliminary examination of invasion patterns revealed that the vast majority of environmental space occupied by the species in their invaded range represented a subset of the conditions occupied in their native range. Consequently, the ‘stability’ and ‘expansion’ values remained near 1 and 0 respectively throughout the time series we analyzed, while ‘unfilling’ declined over time. Accordingly, we defined “spread” as 1 - the ‘unfilling’ metric. Thus, spread in climate space increases over time as a species’ range increases in its invaded territory, occupying a growing subset of the climate space between known to be suitable based on its native distribution (Eckert et al., 2020; Manzoor et al., 2020; Srivastava et al., 2020; Foster et al. 2022).

### Analysis

To determine whether each species’ spread in climate space was best described by a linear, two-phase, or three-phase pattern, we fit their trajectories to linear, asymptotic, and sigmoidal models, respectively (Figure 2). We did this separately for the invaded and native ranges. For the native range, we modelled climate space at 5-year intervals throughout the complete time-series of occurrence records. Both the linear and asymptotic regression models were fit using the R *stats* package (version 3.6.2), using the built-in linear and nonlinear regression fitting functions *lm()* and *SSsymp ()*, respectively. The sigmoidal regression model was fit using *nls()* function, with the maximum and minimum spread values, and the inflection point of the curve (see formula in Figure S2; Ni, 2022).

**Figure 2.**
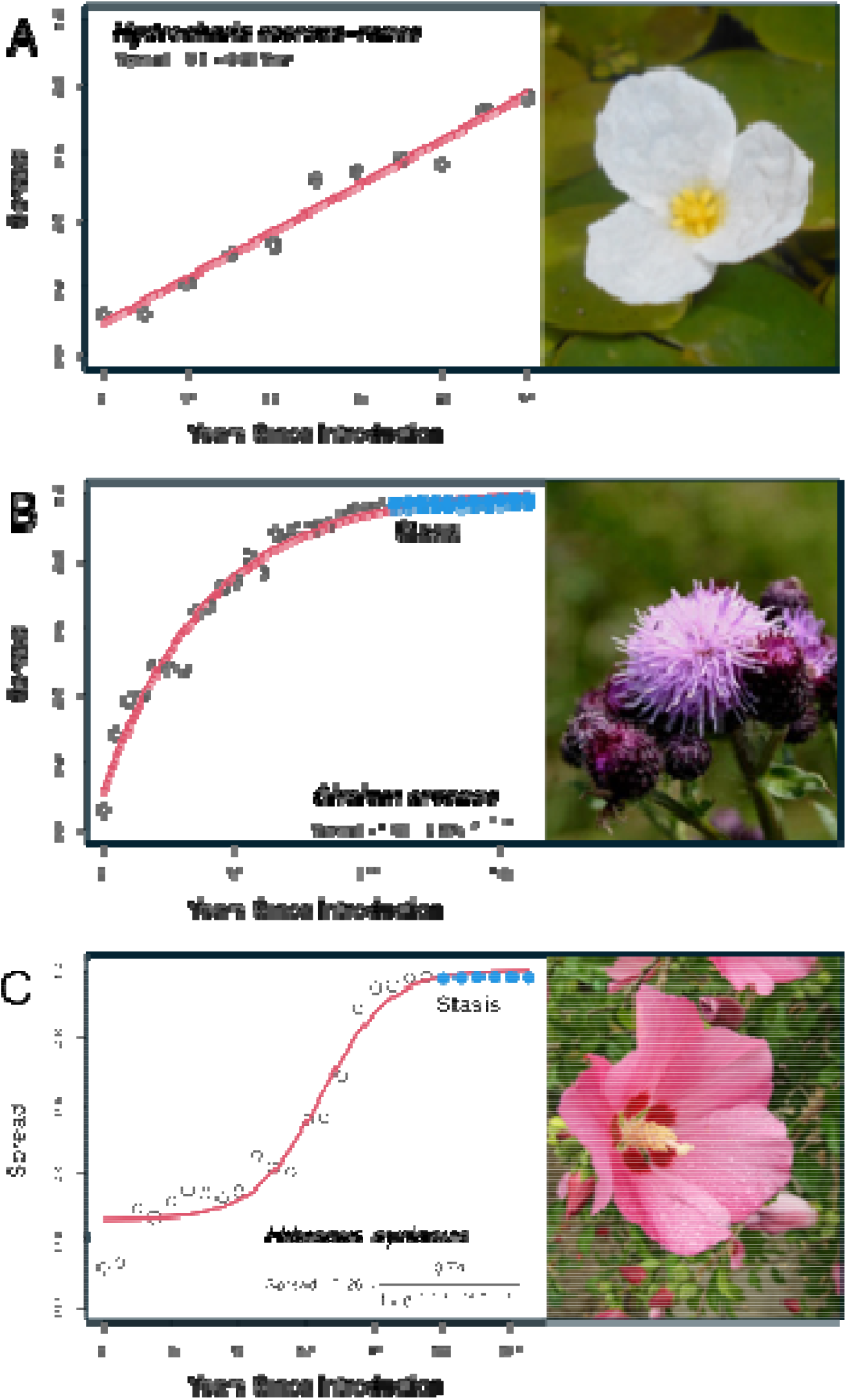
Spread over time plotted for three different species. Spread is calculated as 1 – unfilling; open circles are spread values prior to stasis (i.e., 30 years without increase in spread); closed circles are spread values after stasis; red lines show the best-fit model, with model formulas on the plot. **A.** *Hydrocharis morsus-ranae* shows a single-phase pattern of expansion, best fit by a linear model. **B**. *Cirsium arvense* shows a two-phase pattern of expansion, best fit by an asymptotic model. This species reached environmental stasis (i.e., no further spread) after 105 years. **C**. *Hibiscus syriacus* shows a three-phase expansion, best bit by a sigmoidal model. This species reached environmental stasis after 95 years. Images from Wikipedia Commons.

We compared models using multi-model inference. We used the Akaike Information Criterion (AIC) to determine the model of best fit (i.e., model with lowest AIC). We used the *aictab()* function in R (*stats* package, V3.6.2) to calculate the delta AIC (ΔAIC) among models. To discriminate among competing models, we used a ΔAIC > 2 between the best model and the model with the next smallest AIC (Burnham and Anderson, 2003). When more than one model had a ΔAIC < 2 (i.e., models were indistinguishable), the most parsimonious model was selected (i.e., linear was most parsimonious over asymptotic over sigmoidal).

#### i) Patterns of spread

Species with a single-phase (i.e., linear) pattern of spread were further analyzed to determine if the rate of change was more indicative of a species in active spread (i.e., spreading into climate space at a constant rate over time) or of a species in a lag phase (i.e., climate space occupied remains stable over time). If the rate of change in spread, ß, was less than 0±0.6%/year, the species was classified as being in lag phase, otherwise we classified it as spread. Note that the ecospat calculation of environmental space incorporates both the range of environments occupied and their relative frequency. Consequently, the spread index may decline slightly between time periods as a consequence of shifting proportions, even when the absolute number of observations increases. Accordingly, we allow for slight departures from zero slope (ß) in identifying lag and spread. We set the threshold for ß at 0.6%/year based on visual inspection of the raw data. Finally, we note that these model-based classifications describe statistical patterns in climate space and should not be interpreted as definitive invasion stages.

#### ii) Stasis analysis

For each species, we classified whether they had reached climatic stasis or not in both the invaded and native ranges, and if so, how long it took them to reach it in the invaded range. We defined stasis as a period of no spread for at least 30 years in the last phase of invasion (i.e., excluding any initial lag phase). We selected 30 years as this is the period used to calculate climate normals: a period that it is long enough to account for inter-annual variation without being overly influenced by long-term trends (Arguez et al. 2012). For the stasis analysis, only species previously determined to have a nonlinear spread pattern (i.e., two-phase and three-phase species) were included. Species with a strictly linear spread phase are either still expanding (if ß > 0±0.6 %/year), or still in the lag phase (if ß < 0±0.6).

First, to determine if stasis occurred in the last 30 years (i.e., 1990-2020), we fit a linear model with spread as a function of year for this time period. If the rate of change in spread, ß, was less than 0±0.6 %/year, we categorized this species as having reached stasis. Species with a nonlinear pattern of spread but that did *not* reach stasis before 1990 (i.e., ß > 0±0.6 %/year) were categorized as ‘approaching stasis’.

Second, for species which were in stasis from 1990-2020, we used a backwards 30-year moving window approach to estimate when stasis was first reached. To do this, we shifted the window back in time by 5-year intervals and ran a linear model within each window until ß > 0±0.6 %/year. We used the first year of the earliest 5-year interval of the window to estimate how long it took the species to reach stasis relative to its first recorded occurrence in North America.

Finally, to assess how much unoccupied climate space remains in the introduced range for a species at stasis, we recorded the final spread values for each species determined to have reached stasis.

## Results

Our query generated a set of 1.1 million records spanning 1614 to 2020. Collections increased over time, peaking at 15,945 in 2007. We included 258 species, and 90% of species had more than 1,000 records (range 89 to 17,032, mode 3,698).

### Patterns of spread

Patterns of spread were best fit by a non-linear model for 85.3% (220/258) of the species and by a linear model for the remaining 14.3% (37/258) (Figure 3). For the species best described by a linear model, the climate space occupied remained stable over time for 1.9% (5/258; i.e., lag; slope < |0.6%|) of species, while climate space showed continuous spread (i.e., expanding) for 12.4% (32/258) of species (Figure 3). One species, *Carthamus cretinus*, showed declining environmental occupancy over time. Inspection revealed observations for this species combined an early and apparently short-lived population in South Carolina (five records in 1958-1960), after which all further records (n = 49) were in California. We excluded this species from further analysis due to its idiosyncratic history.

**Figure 3.**
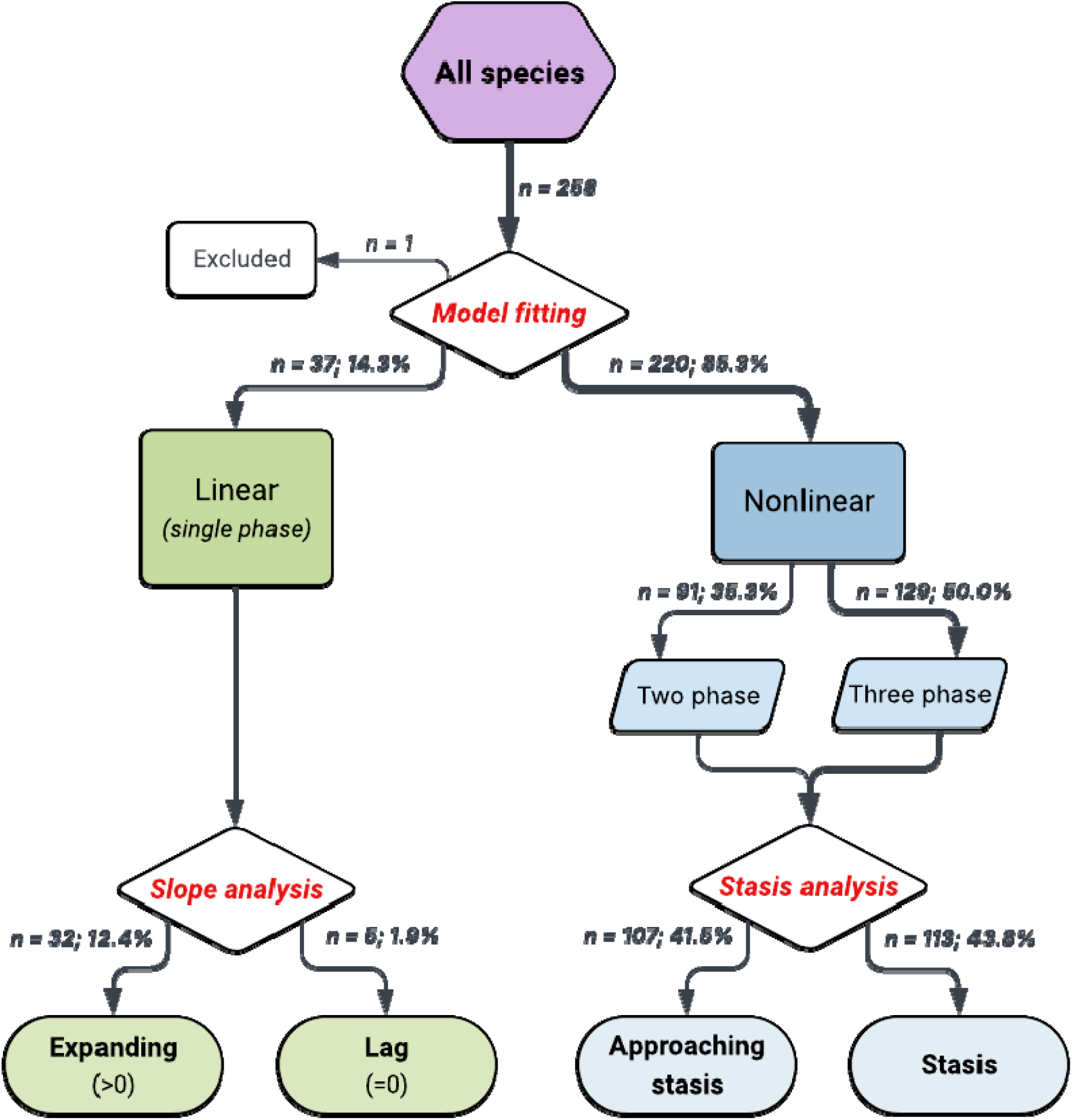
Flow chart summarizing results. Expansion data for all species (n = 258) were input into the model fitting analysis (red), yielding species with a linear pattern over time (green) or with a nonlinear pattern over time (blue). All linear species were classified as ‘single-phase’ and then categorized as ‘expanding’ or ‘lag’. The non-linear species were further classified as either ‘two-phase’ or ‘three-phase’. Both the two and three phase species (all nonlinear) were input into the equilibrium analysis (red), yielding species classified as approaching stasis or having reached stasis. For visualizations of phases, see Figures 1 and 2.

For species with non-linear spread, the pattern of spread was best fit by a two-phase model for 35% (91/258; Figure 3) of species, consistent with early rapid expansion followed by a declining rate of expansion (Figure 2B). The pattern of spread for the rest of the species (58.6%; 129/258) was best fit with a three-phase model (Figure 3) indicating the presence of a lag phase prior to rapid expansion and then a subsequent slowing phase (Figure 3C). Considering all patterns of spread, there was no detectable lag phase in 47.7% (i.e., linear expanding and two-phase expansion, 123/258) of species.

In contrast, the pattern of spread in climate space in the native range of species’ was best explained by a two-phase model for the majority of species (60.5% (156/258), three-phase model for 33.3% (86/258) of species, and the rest were best explained by a linear model (6.2%, 16/258).

### Stasis analysis

There were 113 species (43.8%, n=258) species that reached stasis (i.e., at least 30 years without expansion) and 107 species (41.5%, n=258) that were approaching stasis (Figure 3). In contrast, 218 species (84.5%, n=258) appear to be in climate stasis in their native range. For the species that reached stasis in North America, time to stasis relative to the first recorded occurrence in North America ranged from 30 to 145 years (median: 90 years; Figure 4). When stasis was reached, 97% of suitable climate space was occupied (relative to climate occupied in the native range (Figure S1).

**Figure 4.**
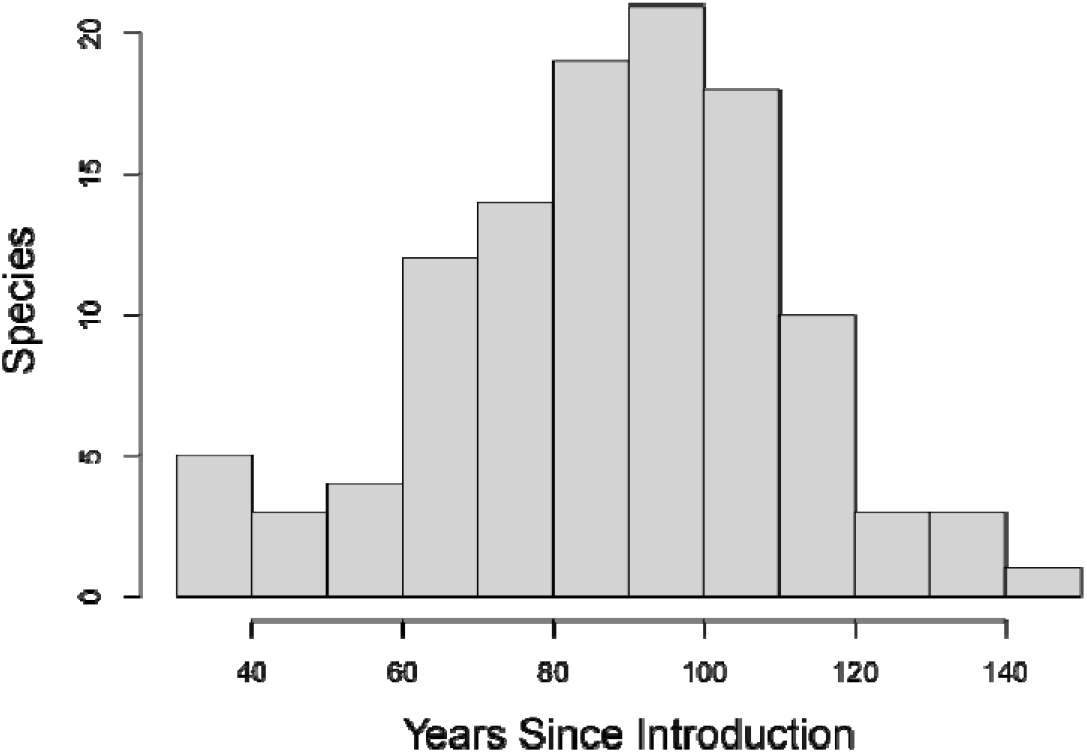
Histogram depicting the distribution of years to stasis across species at stasis (n=113). The mean time to stasis was 90.0 years.

## Discussion

By quantifying the climates occupied by introduced species throughout their invasion history, we used temporal patterns in spread to quantify and describe climatic stasis in North America for more than 250 introduced plants species. Here, we define climatic stasis operationally as a period of at least 30 consecutive years without detectable spread into novel climatic space, based on fitted spread trajectories. This definition provides a consistent temporal criterion for comparing invasion dynamics across species, while acknowledging that climatic stasis does not necessarily imply a fully realized equilibrium distribution. Using this framework, we found that: i) only 43.8% of the species analyzed have reached climatic stasis; ii) it took 90 years on average for a species to reach climatic stasis; iii) species that had reached stasis occupied, on average, 97% of the climates they occupy in their native range in their invaded range; and iv) a detectable lag phase was only present in half of all species (47.7%).

### Climatic stasis

Less than half of all species had reached climatic stasis in North America by 1990, indicating that most of the plants in our study are not yet at equilibrium. While there have been several demonstrations that species distributions are often not at equilibrium (Elith *et al*., 2006; Bradley *et al*., 2015; Briscoe Runquist et al. 2019; Foster et al. 2022), direct comparisons remain difficult because stasis and equilibrium are rarely assessed and definitions vary widely. For example, Osunkoya et al., (2021) reported that only 2% (2/91) of introduced plant species in Australia showed evidence of having reached a stable equilibrium, but they measured equilibrium in geographic space. Ni (2022) found that all plant species they considered displayed a recent slowing of expansion, but they only evaluated the pattern for eight introduced plants and in a much smaller geographical area. Despite these differences, we show that climatic stasis is far less common in invaded ranges (43.8%) than in native ranges (84.5%), supporting the idea that species are much more likely to exhibit stasis in their native distributions (Broennimann and Guisan 2008; Bates and Bertelsmeier 2021).

Consistent with this broader literature, our results suggest that for species that have not yet reached climatic stasis, SDMs constructed to forecast near-future distributions may produce unreliable results. Rare or transient occurrences (waifs) may distort projections (Foster et al. 2022), while assuming all unoccupied space is unsuitable will underestimate potential distributions. For species with a slowing, but not yet stabilized distribution, SDMs may still be useful, but will require caution in their interpretation.

For the species demonstrating climatic stasis, it took 90 years on average to reach it. Comparable equilibrium analyses have typically focused on single species (e.g., Václavík & Meentemeyer, 2012; Briscoe Runquist et al., 2019; Foster et al., 2022) or have evaluated whether equilibrium could be reached without quantifying the time required to do so (e.g., Gallien et al., 2012). Our results highlight how rarely temporal context has been incorporated into equilibrium assessments, despite its conceptual importance.

When stasis *was* reached in the invaded range, species occupied an average of 97% of the climate space they occupy in their native range (Figure S1). This suggests that most species approaching stasis are close to climatic equilibrium as traditionally defined, though stasis alone does not confirm equilibrium. The remaining unoccupied suitable climate space could indicate dispersal barriers that prevent species from reaching all suitable environments, altered biotic interactions or species-environment relationships in the invaded range that reduce their potential distribution relative to the native range (i.e., niche shifts; Mainali et al. 2015). Apparent slowdowns may also arise from changes in sampling intensity through time, saturation of environmental space represented in herbarium collections, or shifts in human-mediated dispersal pathways that decouple spread from climatic suitability. Translating unoccupied climatic space into geographic context will be necessary to assess its biological significance and the likelihood that barriers will be overcome. Nevertheless, the temporal consistency of patterns across species and regions supports the interpretation that these trajectories reflect meaningful components of invasion dynamics.

A few limitations should be considered when interpreting these results. First, estimates of spread dynamics rely on herbarium records, which may incompletely capture early invasion stages or low-density populations, particularly during putative lag phases. Second, our identification of climatic stasis depends on the choice of a 30-year threshold, which, while ecologically motivated, may influence estimates of the proportion of species considered stable. Finally, slowing expansion could in some cases reflect saturation of sampling effort rather than true biological limits. However, the consistency of patterns across hundreds of species suggests that such artefacts are unlikely to fully explain our findings.

### Patterns of spread

We found no detectable lag phase in nearly half of all species, indicating that these species began expanding in environmental space immediately after detection in North America. Previous studies have reported considerable variation in the proportion of plant species exhibiting an initial lag phase. For instance, Osunkoya et al. (2021) observed a lag phase in 59% of 91 introduced plant species in Australia, whereas Hyndman et al. (2015) documented a lag phase in only 28% of introduced plant species in New Zealand and 40% in the Midwestern United States. Such variation across studies is not surprising given that the specific drivers of lag phases are poorly understood. There is evidence that lag times in plants differ across taxonomic groups and native ranges (Larkin, 2012), and that genetic diversity in founding populations and the time required to adapt to novel climatic conditions (i.e., niche shifts) likely contribute to the lag duration, if any (Williamson et al., 2005). Alternatively, lag phases may go undetected if populations remain rare until rapid expansion begins. Regardless, these findings reinforce the importance of early detection and prevention.

We found that a small proportion of species (12%) are currently actively expanding into unoccupied climate space (i.e., linear expansion patterns). On average, these species were introduced to North America 40 years later than species that had non-linear patterns of spread (linear species: mean=1932 ±29 SD; vs. nonlinear species: mean=1892 ±26 SD). This could indicate that species that are currently in an expansion phase have not experienced enough residency time to reach stasis. Osunkoya et al. (2021) reported similar results, with actively expanding species having a time-since-arrival of 37 years less on average than invaders whose expansion had slowed.

Clarifying the factors contributing to variation in invasion trajectories will require a more granular assessment of the traits associated with different invasion patterns as well as the quantification of niche shifts. For example, we would expect species with high dispersal, rapid growth rates, and broad habitat tolerances to require little or no lag period prior to spread. In contrast, traits contributing to density-dependent fitness (i.e., allee effects) such as self-incompatibility, or dependence on specialist pollinators or mycorrhizae could constrain species to a protracted lag period. Past studies have demonstrated clear differences between invasive and non-invasive species in traits related to physiology, shoot allocation, growth rate, size, and fitness (Van Kleunen et al., 2012), but it remains unclear if these traits can help predict how quickly a species reaches stasis. The nature and frequency of a species’ method of introduction, coupled with features of the introduced landscape (i.e., the presence or absence of geographic barriers) could also contribute to the initial rate of spread (Radosevich et al., 2003; Williamson et al., 2005; Larkin, 2012). Similarly, traits such as plant size, phenology and similarity to co-occurring native taxa may contribute to variation in how extensive a newly arrived species needs to be before it is detected and reflected in herbarium collections (Daru et al. 2018). Quantifying the extent of niche shifts that could have occurred in the invaded range could help to disentangle the role of local adaptation and other traits as the factors underlying the variation in the presence—and length—of the lag phase in newly introduced species.

Our results suggest that identifying phases of climatic stasis is valuable in the context of invasive species management and is only possible when spread dynamics are examined through time. Species exhibiting linear expansion likely warrant the highest priority for early detection and rapid response, whereas species with slowing expansion may be more suitable targets for containment or pathway management. Species that have reached climatic stasis, particularly those with a substantial remaining unoccupied climate space (e.g., >15% relative to the native distribution) provide opportunities to investigate the constraints limiting further spread, such as dispersal barriers, biotic interactions or additional environmental factors (e.g., soil or habitat conditions), and to assess whether apparent stability reflects long-term equilibrium or transient barriers. Such periods of stasis or temporary equilibria may therefore provide a window of opportunity for efficient management interventions (Coutts et al., 2018).

### Conclusions

Our study provides a framework to evaluate environmental (in this case climatic) stasis in introduced species using historical observations. We propose environmental stasis is a necessary but not sufficient condition for a species to be at equilibrium distribution. Using this framework, we show that less than half of North American introduced plants analyzed have reached climatic stasis. Future studies should investigate the biological and environmental drivers that determine how quickly species approach stasis. Integrating trait-based, phylogenetic, and genetic data could reveal why some invaders stabilize rapidly while others continue to expand. For those species demonstrating a climatic stasis, mapping unoccupied climate space into geographic context and incorporating non-climatic constraints would help determine whether this is a temporary stasis or potentially a ‘true’ equilibrium. Finally, extending this temporal approach through dynamic SDMs and global comparisons could improve both ecological understanding and the practical forecasting of invasion risks under climate change.

## Data and code availability statement

The data that support the findings of this study are openly available at: https://github.com/plantarum/maisy

## Acknowledgements

We thank Joe Bennett and David Currie for feedback on previous versions of the manuscript. This project was funded by Agriculture and Agri-Food Canada, Project J-002275.

