## Supplementary information for "Invasion histories reveal most North American introduced plants have not yet reached climatic stasis"

Table S1.Breakdown of the native geographic ranges and number of species from each range included in this study.

Figure S1. Distribution of most recent (year 2020) unfilling value for species that reached climatic stasis (n=113).

Figure S2. Overview of the three models used to describe patterns in expansion (*E*) in climate space across species.

**Table S1.** Breakdown of the native geographic ranges and number of species from each range included in this study.

| Native Range | Species |
| --- | --- |
| Asia | 30 |
| Europe | 46 |
| Eurasia | 98 |
| Old World (Europe, Asia and Africa) | 73 |
| South America | 11 |

**Figure S1.** Distribution of most recent (year 2020) unfilling value for species that reached climatic stasis (n=113). Unfilling is the proportion of climate space occupied in the species’ native range that remains unoccupied in the invaded range. The median value is 3.0%.

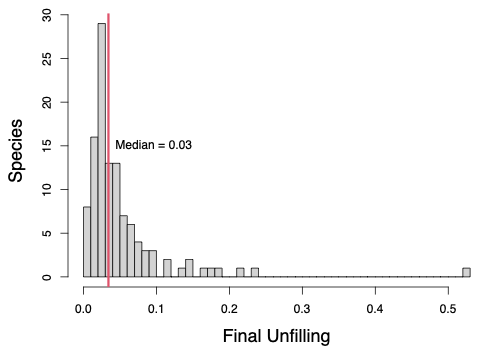

**Figure S2.** Overview of the three models used to describe patterns in expansion (*E*) in climate space across species. Expansion (%) was calculated as 100 – unfilling. Unfilling was calculated as the number of grid cells occupied only in the native range, divided by the total number of cells occupied in the native range. Time is measured in years.

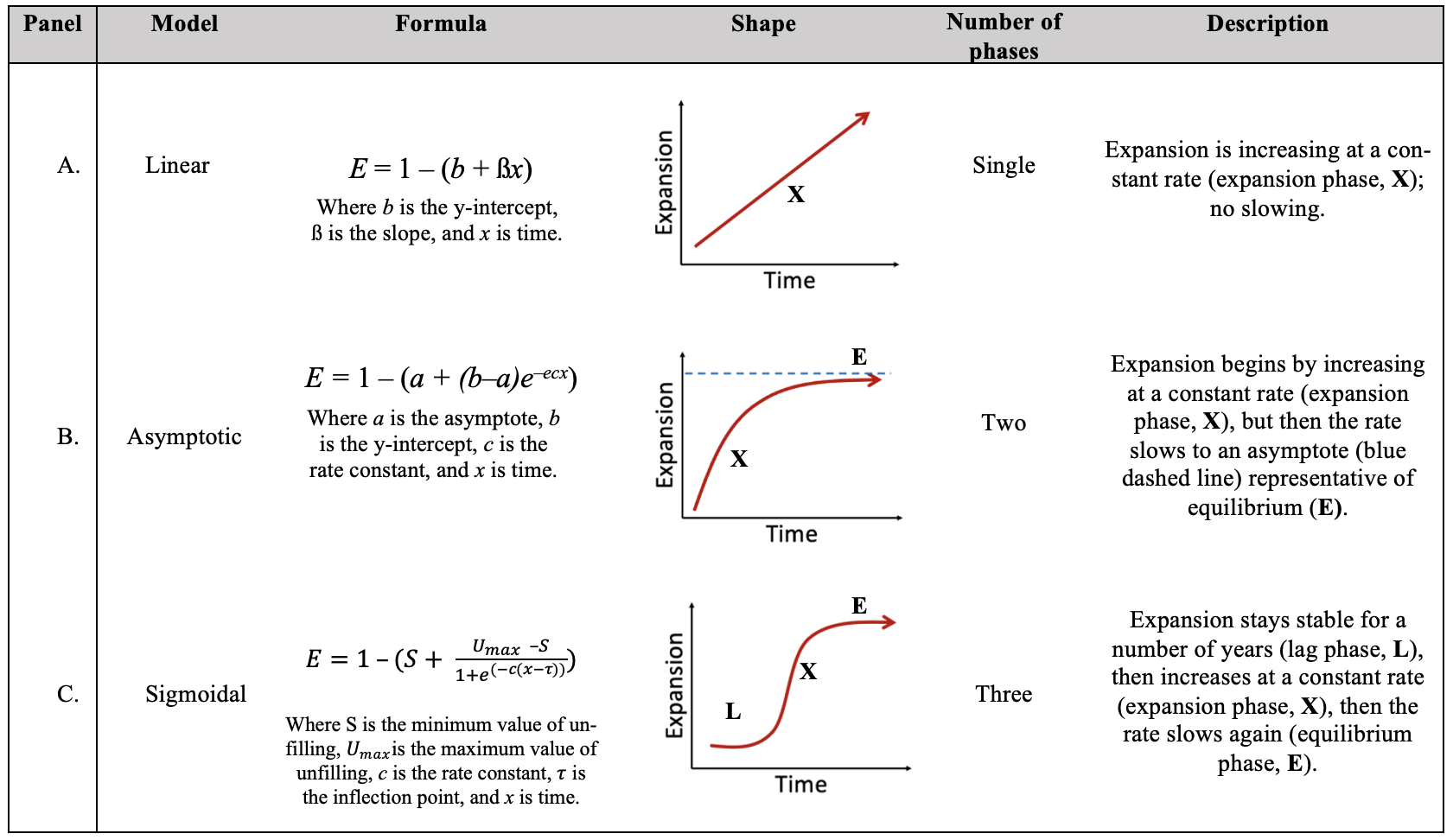
